# Sex-Dependent Age Trajectories of Subcortical Brain Structures: Analysis of Large-Scale Percentile Models and Shape Morphometry

**DOI:** 10.1101/2020.09.30.321711

**Authors:** Christopher R. K. Ching, Zvart Abaryan, Vigneshwaran Santhalingam, Alyssa H. Zhu, Joanna K. Bright, Neda Jahanshad, Paul M. Thompson

## Abstract

Modeling of structural brain variation over the lifespan is important to better understand factors contributing to healthy aging and risk for neurological conditions such as Alzheimer’s disease. Even so, we lack normative data on brain morphometry across the adult lifespan in large, well-powered samples. Here, in a large population-based sample of 26,440 adults from the UK Biobank (age: 44-81 yrs.), we created normative percentile charts for MRI-derived subcortical volumes. Next, we investigated associations between these morphometric measures and the strongest known genetic risk factor for late-onset Alzheimer’s disease (*APOE* genotype) and mapped the spatial distribution of age-by-sex interactions using computational surface mesh modeling and shape analysis. Vertex-wise shape mapping supplements traditional gross volumetric approaches to reveal finer-grained variations across functionally important brain subcompartments. Normative curves revealed volumetric loss with age, as expected, for all subcortical brain structures except for the lateral ventricles, which expanded with age. Surprisingly, no volumetric associations with *APOE* genotype were detected, despite the very large sample size. Age-related trajectories for volumes differed in women versus men, and surface-based statistical maps revealed the spatial distribution of the age-by-sex interaction. Subcortical volumes declined faster in men than women over the full age range, but after age 60, fewer structures showed sex-dependent trajectories, indicating similar volumetric changes in older men and women. Large-scale statistical modeling of age effects on brain structures may drive new insights into individual differences in brain aging and help to identify factors that promote healthy brain aging and risk for disease.

## 1. INTRODUCTION

Regional brain morphometry changes significantly from early development to old age. In neurological and psychiatric disorders, disease-related processes influence brain volumes beyond the changes expected from normal aging. Given the crucial need to distinguish abnormalities in disease from healthy brain variations, a comprehensive statistical model is needed to characterize brain volume variation over the lifespan. Such a model would help establish indicators of both healthy and disordered brain aging, similar to normative growth charts used to track height and weight in childhood (1).

Prior studies of subcortical brain volumes over the lifespan have provided insights but most are limited by modest sample sizes (2). Recent large-scale efforts by the ENIGMA Consortium (3) and the iSTAGING Consortium (4) have pooled datasets from many cohorts worldwide and have dramatically increased sample sizes to improve power and drive consensus findings. Standardized, open-source processing pipelines (such as the gross volume and shape analysis approaches used here) can also improve replication and boost statistical power in neuroimaging studies. The UK Biobank study (5) also offers an opportunity to investigate age effects on brain metrics on an unprecedented scale. In this study, we aimed to study age effects on subcortical brain structures, and the modulating effects of sex and known sources of genetic risk for Alzheimer’s disease. To do this, we applied standardized methods to extract subcortical gross volumes to compute quantile regression models (i.e., percentile charts or nomograms) and model vertex-wise shape variation in a large cohort of adult individuals between 44-81 years of age.

Although regional brain volumes are expected to decline with age, we hypothesized that men and women would differ, on average, in the trajectory of decline. We also expected that carriers of the *APOE* e4 genotype – the strongest known genetic risk factor for late-onset Alzheimer’s disease – would tend to show steeper age-related decline in the hippocampus compared to those carrying *APOE* e3 and e2 genotypes – genotypes associated with typical or lower risk for disease, respectively. We further hypothesized that computational surface modeling combined with 3D statistical shape analysis would reveal complex age, sex and genotypic effects not detected by gross volumetric analysis.

## 2. METHODS

### 2.1 Datasets

T1-weighted brain MRI scans from the UK Biobank (*N*=26,440; age 44-81 years) were collected for an overall sample that was well-balanced for proportions of women and men, and with regard to years of education (**Table 1**).

**Table 1.** Full sample used to generate normative models (nomograms) and for gross volumetric analyses

|  | **Female** | **Male** | **All** |
| --- | --- | --- | --- |
| Mean (SD) | **(N=13,775)** | **(N=12,665)** | **(N=26,440)** |
| **Age (years)** | 62.84 (7.34) | 64.17 (7.62) | 63.5 (7.50) |
| **Education (years)** | 16.31 (3.96) | 16.94 (3.68) | 16.6 (3.84) |

At the time of analysis, 9,414 subjects had available subcortical shape and *APOE* genotype data available (**Table 2**).

**Table 2.** Subsample with available data used for nomograms stratified by *APOE* genotype and for surface-based subcortical shape analyses

|  | **Under Age 60** | **Over Age 60** | **All** |
| --- | --- | --- | --- |
|  | **(N=3,372)** | **(N=6,042)** | **(N=9,414)** |
| **Sex** |  |  |  |
| Female | 1,922 (57.0%) | 2,976 (49.3%) | 4,898 (52.0%) |
| Male | 1,450 (43.0%) | 3,066 (50.7%) | 4,516 (48.0%) |
| **Years of Education (SD)** | 17.09 (3.39) | 16.22 (4.11) | 16.5 (3.89) |
| **Age (SD)** | 54.16 (3.60) | 67.24 (4.27) | 62.56 (7.46) |
| **BMI (SD)** | 26.57 (4.59) | 26.78 (4.11) | 26.7 (4.29) |
| **Genotype** |  |  |  |
| E2E2 | 23 (0.7%) | 40 (0.7%) | 63 (0.7%) |
| E2E3 | 440 (13.0%) | 738 (12.2%) | 1,178 (12.5%) |
| E3E3 | 2,009 (59.6%) | 3,724 (61.6%) | 5,733 (60.9%) |
| E4E3 | 832 (24.7%) | 1,400 (23.2%) | 2,232 (23.7%) |
| E4E4 | 68 (2.0%) | 140 (2.3%) | 208 (2.2%) |

### 2.2 Image Processing Pipelines

Brain MRI scans were segmented using the FreeSurfer software package, version 5.3 (6), with quality control procedures implemented in the ENIGMA consortium (7), to obtain gross volume measures for left and right nucleus accumbens, amygdala, caudate, hippocampus, putamen, pallidum, thalamus and lateral ventricles along with intracranial volume (ICV).

The ENIGMA Shape Analysis pipeline is a surface-based parametric mapping technique used to detect subcortical shape differences across subjects and to map statistical effects of covariates of interest. Using FreeSurfer segmentations as initial inputs, shape registration is based on shape templates and template medial models. Each subcortical structure is represented as triangulated surface mesh. Points on the surface act as vertices that form the overall 3D mesh. A medial model was fit for each structure and was used along with intrinsic shape features to compute a registration to the template (8, 9).

Two point-wise measures of shape morphometry were derived. The first, termed radial distance or thickness, was calculated using the medial model approach where for each point *p* ∈ *M* on the surface, and given a medial curve *c*: [0,1] → *R*^3^. The radial distance is defined by:

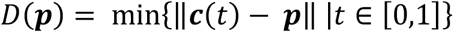

The second measure, based on surface Tensor Based Morphometry (TBM), generalizes TBM on Euclidean spaces to surfaces (9). The differential map between the tangent spaces of two surfaces replaces the Jacobian:

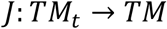

In our model, *M_t_* is the average template, and *M* is the surface we wish to study. *J* is a linear mapping, and may be thought of as the restriction of the standard Jacobian determinant to the tangent spaces of the template and study surfaces. While analysis of the full tensor using log-Euclidean metrics on symmetric positive-definite matrices is possible (9, 10), these analyses are difficult to interpret. Our model instead considers the Jacobian determinant, representing the surface dilation ratio between the template and the study subject. An interpretation of this measure is the areal dilation or contraction required to match a small surface patch around a particular point of the subject surface to the corresponding point on the template. A higher Jacobian indicates a larger volume of a structure’s subfield corresponding to the region. Our final TBM measure is the logarithm of the Jacobian determinant, to obtain a distribution closer to Gaussian.

In this way, both radial distance (termed *thickness* from now on) and the logarithm of the Jacobian determinant (termed simply *Jacobian* from now on) were calculated in native space for up to 2,502 points across each subcortical structure, providing an index of regional shape differences across subjects (**Figure 1**). The ENIGMA Shape Analysis pipeline has been applied in several recent large-scale studies mapping subcortical effects of substance abuse (11), major depression (12) and 22q11.2 deletion syndrome (13).

**Figure 1.**
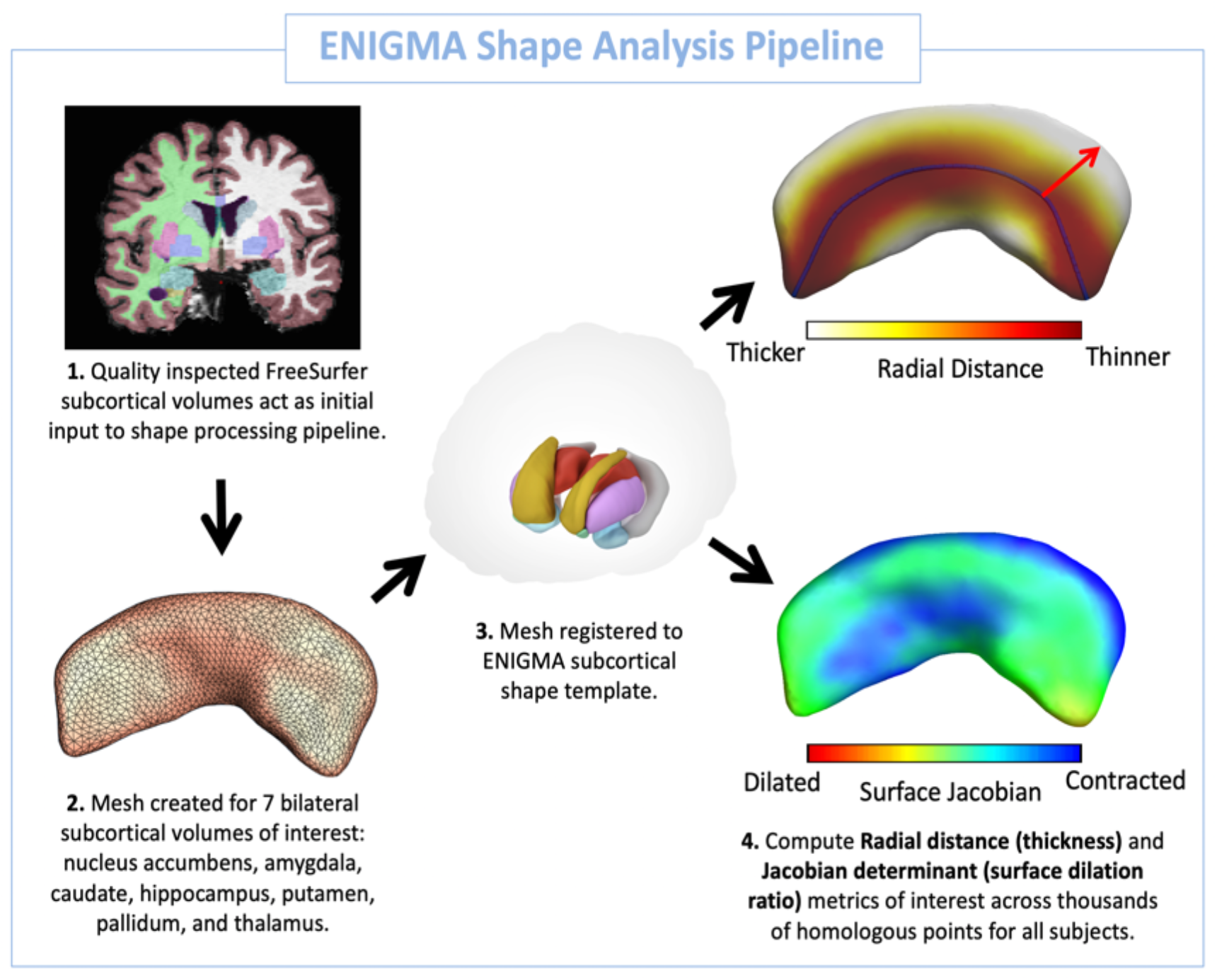
ENIGMA Shape Analysis Pipeline (http://enigma.ini.usc.edu/ongoing/enigma-shape-analysis/). Shape models are generated for the caudate, putamen, pallidum, hippocampus, amygdala, thalamus and nucleus accumbens.

### 2.3 Modeling and Statistical Analysis

The Python class *LinearRegression* from the *scikit-learn* library was used to adjust the subcortical volumes (average of left and right structures) for linear effects of ICV prior to fitting quantile regression models (termed *nomograms*, from now on). The Python library *statsmodels*, specifically class *quantreg*, was used to compute percentile curves for each structure based on standardized residuals, with structure volume as the dependent variable and age as the independent variable. We fit separate models for men and women and for all *APOE* genotypes to visualize age trends, stratified by genotype and by sex.

Statistical analysis involved the application of multiple linear regression (*lm* package in R) to model the association between age, sex, age-by-sex interactions, and *APOE* genotype with both gross volumes and shape metrics (Jacobian and thickness) while adjusting for age, ICV, years of education, body mass index, and adjusting for multiple comparisons (FDR correction *q*<0.05).

## 3. RESULTS

As expected, nomograms showed decreasing volume trends with age, for all structures, except for the lateral ventricles, whose volumes increased with age (**Figure 2**). A significant age-by-sex interaction in all gross volumes showed greater volumetric changes with age in men compared to women across the full age range (**Table 3**). However, in individuals over 60 years old, this age-by-sex interaction was attenuated with fewer structures showing differential trajectories, indicating similar volumetric changes in older men and women. Shape analysis revealed complex age-by-sex effects, with subregions of certain structures showing both higher and lower rates of thickness and surface area loss in men compared to women with increasing age (**Figure 4**).

**Figure 2.**
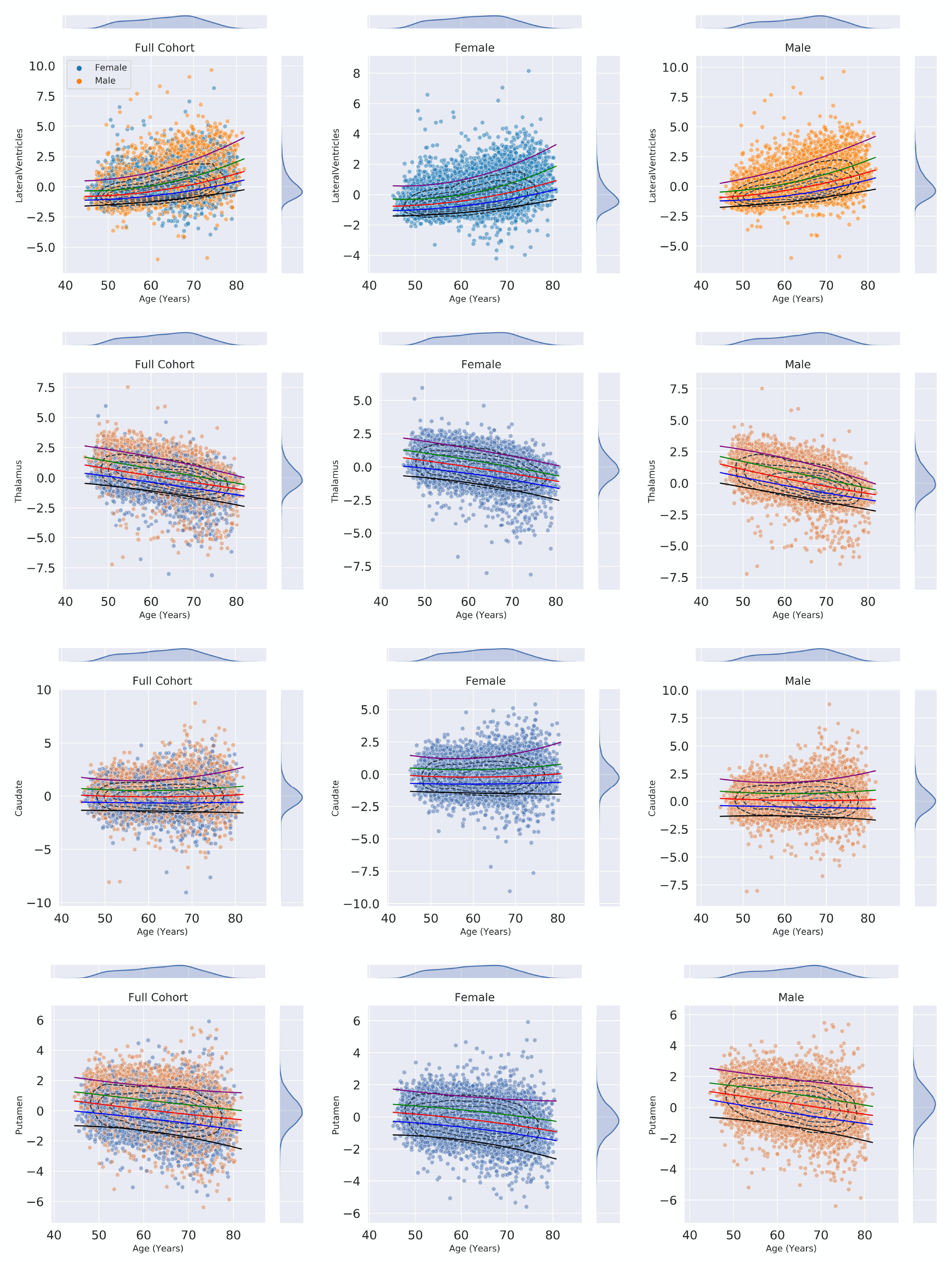
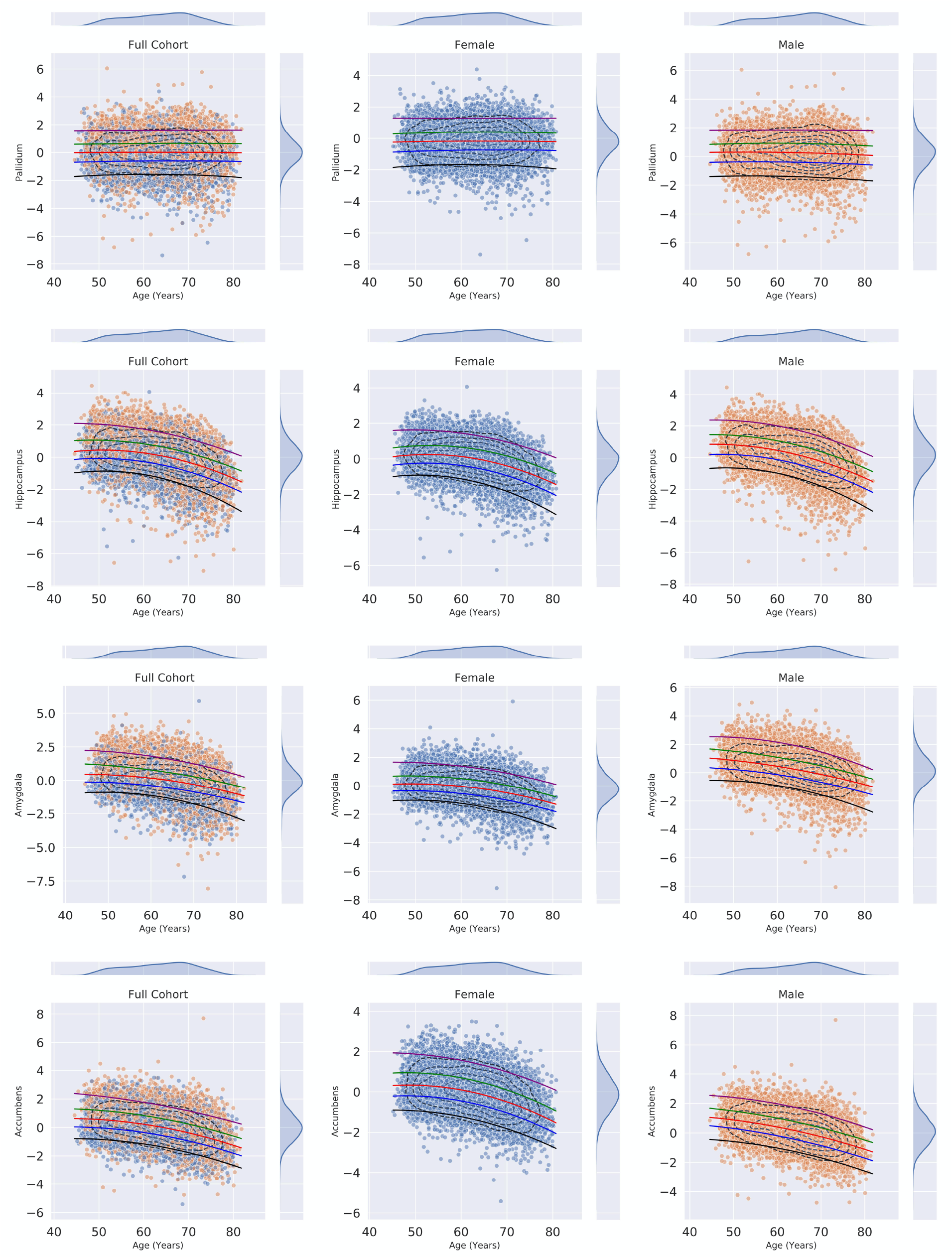
Nomograms for average left and right gross subcortical volumes in the full cohort (N=26,440). *Left column:* full cohort; *middle column:* females; *right column:* males.

No effect of *APOE* genotype nor any interaction between sex, age and genotype were observed, possibly due to the small, underpowered samples of *APOE* e2/2 and e4/4 genotypes (**Table 3 and Figure 3**).

**Table 3.** Statistical analyses of left and right gross volumes in the full cohort. A. **Effect of age** adjusted for age^2^, sex, years of education, BMI and ICV. B. **Effect of sex** adjusted for age, age^2^, years of education, BMI and ICV. C. **Age-by-sex interaction** adjusted for age, age^2^, sex, years of education, BMI, and ICV. D. **Age-by-sex interaction in individuals >60 years** of age adjusted for age, age^2^, sex, years of education, BMI, and ICV. Highlighted cells indicate significant results after correction for multiple comparisons (FDR *q*<0.05). L:left; R:right; LatVent: lateral ventricles; thal: thalamus; caud: caudate; put: putamen; pal: pallidum; hippo: hippocampus; amyg: amygdala; accumb: accumbens; ICV: intracranial volume.

| A | ROI | beta | p |
| --- | --- | --- | --- |
|  | LLatVent* | 9.75 | 3.28E-21 |
|  | RLatVent* | 9.88 | 1.78E-24 |
|  | Lthal | -0.21 | 1.29E-01 |
|  | Rthal | -0.08 | 4.21E-01 |
|  | Lcaud* | 0.24 | 1.18E-03 |
|  | Rcaud* | 0.16 | 3.53E-02 |
|  | Lput | -0.03 | 8.10E-01 |
|  | Rput | -0.09 | 3.41E-01 |
|  | Lpal* | 0.13 | 1.52E-03 |
|  | Rpal | -0.02 | 4.88E-01 |
|  | Lhippo* | -0.85 | 1.73E-31 |
|  | Rhippo* | -0.94 | 2.30E-38 |
|  | Lamyg* | -0.27 | 1.84E-14 |
|  | Ramyg* | -0.29 | 7.31E-17 |
|  | Laccumb* | -0.13 | 2.77E-13 |
|  | Raccumb* | -0.12 | 1.47E-13 |
|  | ICV* | 57.42 | 2.36E-02 |

| B | ROI | beta | p |
| --- | --- | --- | --- |
|  | LLatVent* | 1270.74 | 3.03E-21 |
|  | RLatVent* | 1078.50 | 1.10E-17 |
|  | Lthal* | 291.92 | 3.91E-59 |
|  | Rthal* | 250.69 | 3.29E-87 |
|  | Lcaud* | 91.84 | 1.42E-21 |
|  | Rcaud* | 110.80 | 7.93E-30 |
|  | Lput* | 271.11 | 1.71E-86 |
|  | Rput* | 296.73 | 1.66E-135 |
|  | Lpal* | 82.36 | 2.91E-55 |
|  | Rpal* | 77.52 | 3.07E-86 |
|  | Lhippo* | 156.86 | 1.27E-61 |
|  | Rhippo* | 146.46 | 4.09E-54 |
|  | Lamyg* | 115.27 | 1.22E-134 |
|  | Ramyg* | 127.24 | 2.72E-166 |
|  | Laccumb* | 25.94 | 3.19E-29 |
|  | Raccumb* | 33.82 | 9.06E-54 |
|  | ICV* | 152913.66 | 2.00E-16 |

| C | ROI | beta | p |
| --- | --- | --- | --- |
|  | LLatVent* | 131.34 | 1.77E-16 |
|  | RLatVent* | 112.65 | 5.14E-14 |
|  | Lthal* | -10.38 | 1.11E-06 |
|  | Rthal* | -9.05 | 1.37E-09 |
|  | Lcaud* | -3.81 | 8.81E-04 |
|  | Rcaud* | -3.24 | 5.29E-03 |
|  | Lput* | -5.33 | 1.02E-03 |
|  | Rput* | -4.96 | 4.20E-04 |
|  | Lpal | -1.29 | 3.87E-02 |
|  | Rpal* | -0.31 | 5.00E-01 |
|  | Lhippo* | -8.43 | 5.13E-14 |
|  | Rhippo* | -7.43 | 3.22E-11 |
|  | Lamyg* | -2.78 | 3.67E-07 |
|  | Ramyg* | -2.59 | 1.75E-06 |
|  | Laccumb* | -1.18 | 1.84E-05 |
|  | Raccumb* | -0.62 | 1.77E-02 |
|  | ICV* | 1016.50 | 9.83E-03 |

| D | ROI | beta | p |
| --- | --- | --- | --- |
|  | LLatVent* | 129.82 | 6.78E-04 |
|  | RLatVent* | 98.80 | 5.59E-03 |
|  | Lthal | -3.00 | 5.09E-01 |
|  | Rthal | -1.50 | 6.44E-01 |
|  | Lcaud* | -10.64 | 5.13E-05 |
|  | Rcaud* | -9.68 | 2.80E-04 |
|  | Lput | -2.45 | 4.96E-01 |
|  | Rput | -1.02 | 7.48E-01 |
|  | Lpal | -1.18 | 3.82E-01 |
|  | Rpal | -0.53 | 6.13E-01 |
|  | Lhippo | -4.46 | 7.51E-02 |
|  | Rhippo | -2.70 | 2.81E-01 |
|  | Lamyg | -0.80 | 5.11E-01 |
|  | Ramyg | -0.05 | 9.67E-01 |
|  | Laccumb | -0.33 | 5.88E-01 |
|  | Raccumb | 0.21 | 7.19E-01 |
|  | ICV | -300.88 | 7.43E-01 |

**Figure 3.**
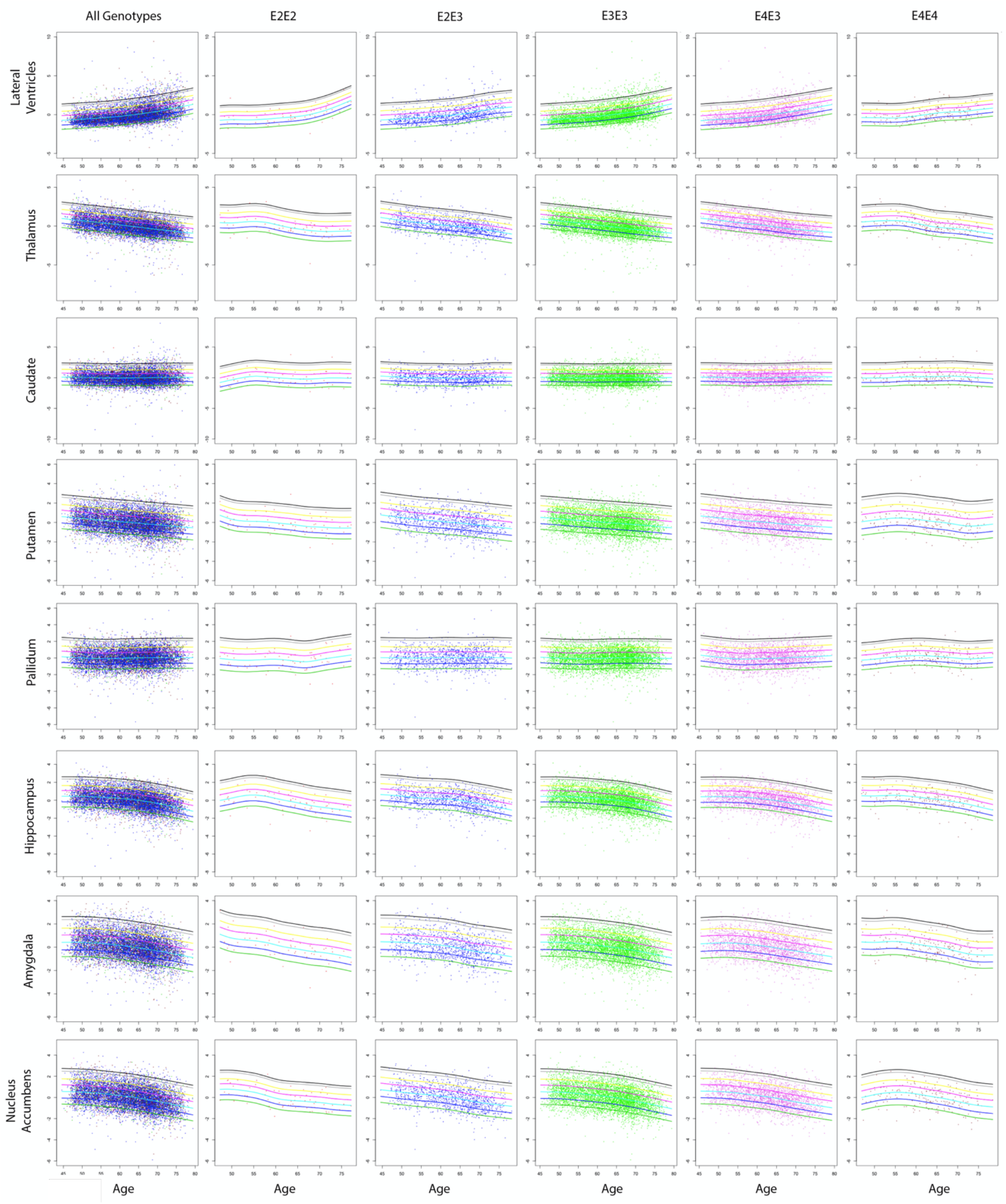
Nomograms for average left and right gross subcortical volumes, in the full sample, and in groups split by *APOE* genotype. All structures show a volume decline with age, in the age range studied here, except for the lateral ventricles, which expand with age (*top row*). Age trends stratified by genotype are shown in columns 2-6; these trends are statistically compared using formal tests, reported in the main text. Percentile curves included for 99%, 98%, 90%, 75%, 50%, 25% and 10%.

**Figure 4.**
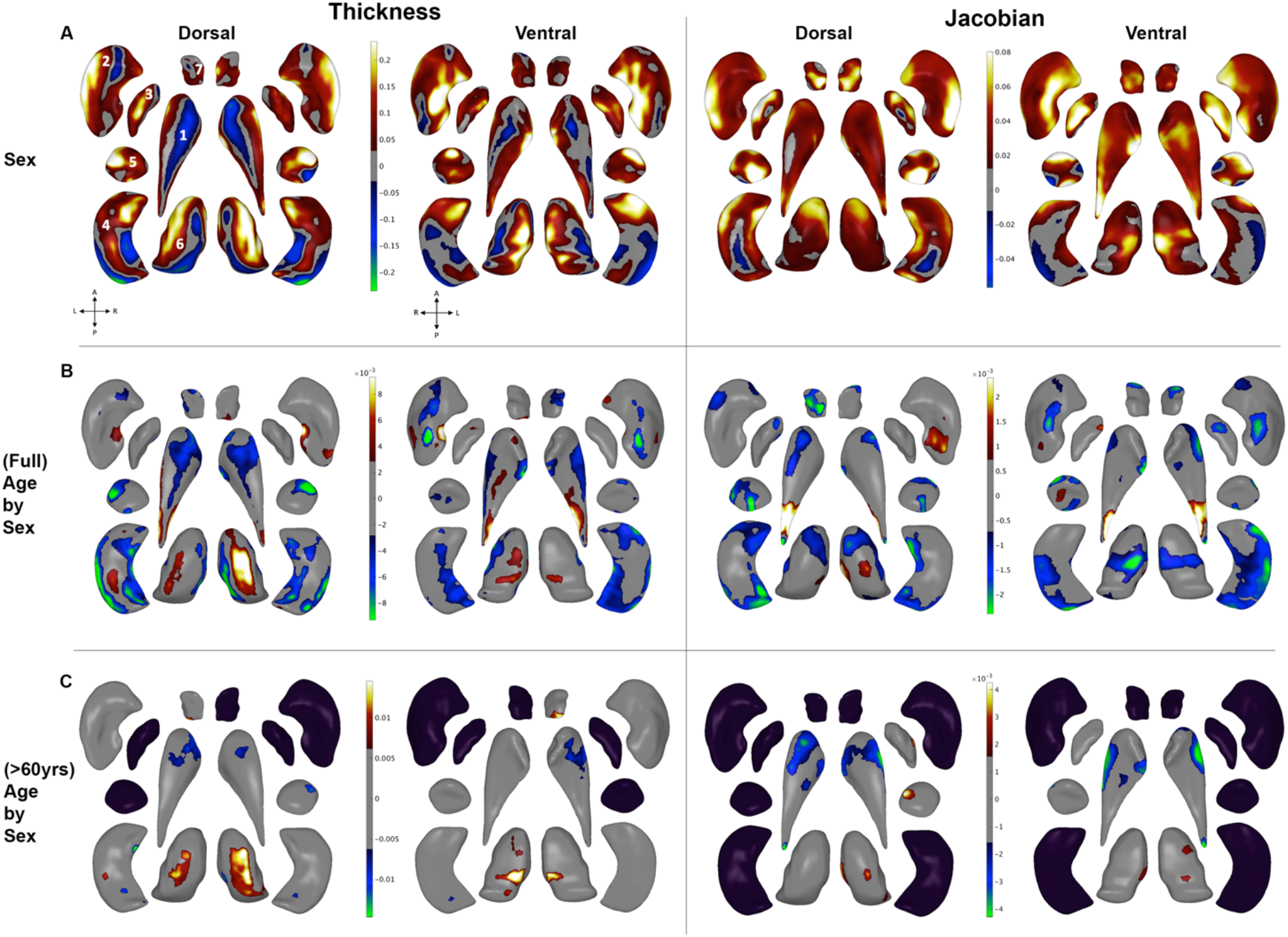
Shape analysis of subcortical brain structure, with regression coefficients plotted in regions passing correction for multiple comparisons (FDR *q*<0.05). A. **Effect of sex** on thickness and Jacobian adjusted for age, age^2^, years of education, BMI and ICV. Warm (*red/yellow*) colors indicate positive model coefficients, or regions of greater Jacobian (surface area) or thickness values in men compared to women. Cooler (*blue/green*) colors indicate negative model coefficients, or regions of lower Jacobian (surface area) or thickness measures in men compared to women. B. **Age-by-sex interaction** adjusted for age, age^2^, sex, years of education, BMI and ICV. Blue/green regions indicate greater local surface area or thickness decline in men compared to women with increasing age. C. **Age-by-sex interaction in individuals >60 years of age** adjusted for age, age^2^, sex, years of education, BMI and ICV. 1: caudate; 2: putamen; 3: pallidum; 4: hippocampus; 5: amygdala; 6: thalamus; 7: nucleus accumbens. Gray regions indicate non-significant vertices after correction for multiple comparisons. Black structures are those for which no vertex-wise test was significant after correction for multiple comparisons.

## 4. DISCUSSION

Here, in one of the largest subcortical brain morphometry studies to date, we provide five key findings. First, nomograms indicate generally decreasing volume trends for all structures over the adult age range studied, except for the lateral ventricles whose volumes increased with age, as expected. Second, differential aging trajectories for women and men, as visualized in the nomograms from the full cohort (N=26,440), were confirmed with formal statistical analysis of gross volumes and with shape analyses. A significant age-by-sex interaction in all gross volumes showed greater volumetric changes with age in men than women across the full age range studied. Third, in individuals over 60 years old, an attenuated age-by-sex interaction was observed, with fewer structures showing significant differential trajectories in men versus women, indicating similar volumetric changes in older men and women. Fourth, shape analysis revealed patterns of complex age-by-sex effects over the full age range, with subregions of certain structures showing both higher and lower rates of thickness and surface area loss in men compared to women with increasing age. Again, in individuals over 60 years old, that interaction was attenuated with shape maps revealing fewer vertices with a significant age-by-sex interaction. Fifth, no significant association with *APOE* genotype was detected in the current sample, though nomograms show the need for more study samples to better model age effects in groups carrying at least one *APOE* e2 or *APOE* e4 allele.

The failure to detect an *APOE* effect may be cohort specific, as previous UK Biobank analyses have not detected the hypothesized *APOE* effects (14). The worldwide frequency of *APOE* e2, e3 and e4 alleles are 8.4%, 77.9% and 13.7%, respectively (15). Even large-scale, population-based samples such as UK Biobank struggle to collect data from enough e2 and e4 carriers to empower analyses of the documented protective and detrimental effects of these genotypes in the general population. As the UK Biobank study continues to release more MRI data, our ability to model the normative trajectories of these less common but highly impactful genetic factors may improve.

The current study has several limitations. First, although we refer to the age effects as trajectories, the pattern of aging is inferred here from cross-sectional data; trajectories fitted from longitudinal data may better reflect the changes occurring for specific individuals in the population. Especially at older ages, poorer health, mortality, and attrition effects may limit the individuals represented in the study, so the brain metrics in older ages may represent people in better health for their age than in the younger portion of the age range. Applying these nomograms to individuals beyond the UK biobank would require adaptation. As different scanners and imaging protocols differ in their signal to noise ratio for the measures examined here, the models provided here may not extend to data collected from different scanners or using different acquisition protocols. Methods to adapt nomograms and statistics across cohorts are under rapid development and include ComBat, its particular variants (ComBat-GAM, CovBat, and longitudinal ComBat) and hierarchical Bayesian methods in which the mean, variance, and other moments of the data distribution can be modeled as arising from a random process whose distributional parameters are also estimated. Such harmonization efforts will allow nomogram data to inform the analysis of new datasets, as well as assist in the cooperative analysis of both small and large datasets for specific questions.

In summary, we presented aging trajectories for subcortical brain volumes and were able to map a spatially complex age-by-sex interaction using high-resolution shape morphometry. Ongoing work aims to more closely investigate the differential aging effects in men and women that map on to underlying subfields or nuclei with known structural and functional brain connectivity. Analyses incorporating measures of white matter integrity from the same UK Biobank sample will help identify potential structural vulnerabilities that could inform future mechanistic insights into healthy aging and age-related brain disorders such as Alzheimer’s disease.

## 5. ACKNOWLEDGMENTS

CRKC, PMT, and SIT were supported in part by NIH Grant U54 EB020403, R01 MH116147, R56 AG058854, P41 EB015922, and R01MH111671; CRK was supported by NIA T32AG058507. CRKC, NJ, and PMT received partial research support from Biogen, Inc. (Boston, USA).

## Notes

### Competing Interest Statement

The authors have declared no competing interest.

## REFERENCES

[1] World Health Organization. Nutrition for Health and Development., “WHO child growth standards: growth velocity based on weight, length and head circumference: methods and development,” Geneva, Switzerland: World Health Organization, Department of Nutrition for Health and Development, (2009).

[2] Tullo, S., Patel, R., Devenyi, G. A., Salaciak, A., Bedford, S. A., Farzin, S. et al., “MR-based age-related effects on the striatum, globus pallidus, and thalamus in healthy individuals across the adult lifespan,” Hum Brain Mapp, vol. 40, pp. 5269–5288 (2019).

[3] Dima, D., Papachristou, E., Modabbernia, A., Doucet, G. E., Agartz, I., Aghajani, M. et al. “Subcortical Volume Trajectories across the Lifespan: Data from 18,605 healthy individuals aged 3-90 years” BioRxiv. doi: https://doi.org/10.1101/2020.05.05.079475 (2020).

[4] Pomponio, R., Erus, G., Habes, M., Doshi, J., Srinivasan, D., Mamourian, E., et al., “Harmonization of large MRI datasets for the analysis of brain imaging patterns throughout the lifespan,” Neuroimage, vol. 208, p. 116450 (2020).

[5] Miller, K. L., Alfaro-Almagro, F., Bangerter, N. K., Thomas, D. L., Yacoub, E., Xu, J., et al., “Multimodal population brain imaging in the UK Biobank prospective epidemiological study,” NatNeurosci, vol. 19, pp. 1523–1536 (2016).

[6] Fischl, B., Salat, D. H., Busa, E., Albert, M., Dieterich, M., Haselgrove, C., et al., “Whole brain segmentation: automated labeling of neuroanatomical structures in the human brain,” Neuron, vol. 33, pp. 341–55 (2002).

[7] http://enigma.ini.usc.edu/protocols/imaging-protocols/

[8] Gutman, B. A., Fletcher, P. T., Cardoso, M. J., Fleishman, G. M., Lorenzi, M., Thompson, P. M., et al., “A Riemannian Framework for Intrinsic Comparison of Closed Genus-Zero Shapes,” Inf Process Med Imaging, vol. 24, pp. 205–18 (2015).

[9] Gutman, B. A., Jahanshad, N., Ching, C. R., Wang, Y., Kochunov, P. V., Nichols, T. E., et al., “Medial Demons Registration Localizes The Degree of Genetic Influence Over Subcortical Shape Variability: An N= 1480 Meta-Analysis,” Proc IEEE Int Symp Biomed Imaging, vol. 2015, pp. 1402–1406 (2015).

[10] Wang, Y., Song, Y., Rajagopalan, P., An, T., Liu, K., Chou, Y. Y., et al., “Surface-based TBM boosts power to detect disease effects on the brain: an N=804 ADNI study,” Neuroimage, vol. 56, pp. 1993–2010 (2011).

[11] Chye, Y., Mackey, S., Gutman, B. A., Ching, C. R. K., Batalla, A., Blaine, S., et al., “Subcortical surface morphometry in substance dependence: An ENIGMA addiction working group study,” Addict Biol, p. e12830 (2019).

[12] Ho, T. C., Gutman, B., Pozzi, E., Grabe, H. J., Hosten, N., Wittfeld, K., et al., “Subcortical shape alterations in major depressive disorder: Findings from the ENIGMA major depressive disorder working group,” Hum Brain Mapp (2020).

[13] Ching, C. R. K., Gutman, B. A., Sun, D., Villalon Reina, J., Ragothaman, A., Isaev, D., et al., “Mapping Subcortical Brain Alterations in 22q11.2 Deletion Syndrome: Effects of Deletion Size and Convergence With Idiopathic Neuropsychiatric Illness,” Am J Psychiatry, vol. 177, pp. 589–600 (2020).

[14] Lyall, D. M., Ward, J., Ritchie, S. J., Davies, G., Cullen, B., Celis, C., et al., “Alzheimer disease genetic risk factor APOE e4 and cognitive abilities in 111,739 UK Biobank participants,” Age Ageing, vol. 45, pp. 511–7 (2016).

[15] Liu, C. C., Kanekiyo, T., Xu, H., Bu, H. “Apolipoprotein E and Alzheimer disease: risk, mechanisms and therapy,” Nat Rev Neurol, vol. 9, pp. 106–18 (2013).

